# Clathrin light chains CLCa and CLCb have non-redundant roles in epithelial lumen formation

**DOI:** 10.1101/2023.04.25.538235

**Authors:** Yu Chen, Kit Briant, Marine D. Camus, Frances M. Brodsky

## Abstract

To identify functional differences between vertebrate clathrin light chains (CLCa or CLCb), phenotypes of mice lacking genes encoding either isoform were characterised. Mice without CLCa displayed 50% neonatal mortality, reduced body weight, reduced fertility, and ∼40% of aged females developed uterine pyometra. Mice lacking CLCb displayed a less severe weight reduction phenotype compared to those lacking CLCa, and had no survival or reproductive system defects. Analysis of female mice lacking CLCa that developed pyometra revealed ectopic expression of epithelial differentiation markers (FOXA2 and K14) and a reduced number of endometrial glands, indicating defects in the lumenal epithelium. Defects in lumen formation and polarity of epithelial cysts derived from uterine or gut cell lines were also observed when either CLCa or CLCb were depleted, with more severe effects from CLCa depletion. In cysts, the CLC isoforms had different distributions relative to each other, while they converge in tissue. Together, these findings suggest differential and cooperative roles for CLC isoforms in epithelial lumen formation, with a dominant function for CLCa.

## Introduction

Clathrin-mediated membrane traffic is critical for a range of biological processes including tissue development, neurotransmission, metabolism and immunity (1,2,3). Clathrin-mediated endocytosis and recycling from endosomes are responsible for regulating plasma membrane levels of numerous receptors and transporters, while clathrin-mediated transport at intracellular membranes influences formation of lysosomes, secretory granules and specialised organelles (2,4). Adaptor recruitment to specific membranes leads to localised clathrin self-assembly, deforming the membrane and selectively capturing cargo into clathrin-coated vesicles (CCVs) for ongoing transport to target destinations (1,5). The major form of clathrin in all eukaryotes is a triskelion-shaped trimer of three identical clathrin heavy chain subunits (CHC17 in humans) with three associated clathrin light chains (CLCs). In some vertebrate species (present in humans, absent from rodents and ruminants), there is a second clathrin formed by CHC22 that does not bind CLCs, which is muscle-enriched and mediates specialised membrane traffic of the GLUT4 glucose transporter (1,2). In all vertebrates, the CLCs are obligate subunits for the ubiquitous CHC17 clathrin and are encoded by two genes that undergo tissue-specific splicing, respectively producing CLCa and CLCb proteins, each of which has a ubiquitously expressed or neuron-specific splice variant (denoted by the prefix ‘u’ and ‘n’ respectively) (6,7,8). uCLCa and uCLCb are ∼60% identical in protein sequence with these differences, as well as their tissue-specific expression levels and splicing patterns highly conserved across vertebrates (6,9). These characteristics suggest evolutionary pressure to maintain the different CLC isoforms, suggesting the CLCs perform non-redundant and tissue-specific physiological functions. Here we investigate isoform-specific CLC functions in mice following homozygous deletion of CLC-encoding genes (*Clta* and *Cltb*).

Since their identification as components of the clathrin triskelion in 1980 and appreciation of their diversity in vertebrates (10,11), it has been challenging to discover differential roles for the CLCs. The only completely shared domain between CLCa and CLCb is a 22 amino acid consensus sequence (about 10% total length) close to their N-termini that is responsible for binding huntingtin-interacting proteins (HIP1 and HIP1R), with a homologous sequence binding the HIP-related Sla2p in yeast (12,13,14). The three C-terminal residues of the consensus sequence are essential for CLC’s role in modulating the pH dependence of clathrin self-assembly (15). Other than the consensus sequence, the N-terminal thirds of CLCa and CLCb are divergent in length and sequence, while they are more similar in the central CHC-binding domain (66% identity) and C-terminal third (76% identity) with neuronal splicing inserts at equivalent positions (7,8). nCLCa includes 30 amino acids encoded by two exons and nCLCb includes 18 amino acids encoded by one exon with homology to the first 18 residues of the nCLCa insert (7). Functional studies of CLC contribution to clathrin pathways in tissue culture systems have revealed roles in G-protein-coupled receptor (GPCR) uptake, focal adhesion formation, cell migration, and invadopodia formation (16,17,18,19,20). In only a few of these have isoform-specific properties of CLCs been identified. The epithelial splice variant of myosin VI was shown to interact with clathrin specifically through CLCa, with consequences for CME at the actin-rich apical membrane of these cells (21), and a preferential role for CLCa was identified in cell spreading and migration (19), while isoform-specific CLCb phosphorylation was shown to influence clathrin dynamics and GPCR uptake (16). In complementary *in vitro* studies, reconstitution of clathrin with different CLC isoforms showed isoform-specific effects on biophysical properties of the clathrin lattice (22). Thus, how CLCs diversify clathrin function to meet specialised, tissue-specific needs *in vivo* remains a key question in the field. To this end, we generated homozygous mice lacking either *Clta* or *Cltb*. In these animals, we previously characterised a role for CLCa in membrane traffic controlling antibody isotype switching in B lymphocytes, which express mainly CLCa (9). We also found that loss of nCLCa and nCLCb has opposing effects on synaptic vesicle generation and synaptic transmission (22). Notably, CLCa loss reduced synaptic vesicle formation and transmission, while both were increased in CLCb-depleted mice (22). These neuronal phenotypes suggested that CLCa performs more of a housekeeping role for clathrin, while CLCb acts to attenuate or regulate CLCa function, perhaps through its natural competition with CLCa for CHC binding (23), such that a balance of the two CLC isoforms is required for normal function.

Here, we further characterised the CLCa and CLCb knock-out (KO) mice to elucidate the physiological roles of CLCa and CLCb. We found that CLCa KO mice have much stronger phenotypes than CLCb. Approximately half of CLCa KO mice die within three days of birth. Surviving CLCa KO mice have reduced bodyweight and impaired fertility. In contrast, CLCb KO mice show very mild phenotypes, with no mortality or infertility associated with this genotype, and only a small reduction in bodyweight. Furthermore, we observed that loss of CLCa, but not CLCb, in female mice results in a change in endometrial epithelial cell identity and the development of uterine inflammation, indicating a critical isoform-specific role for CLCa in these cells. When analysed *in vitro*, we found that acute loss of either CLCa or CLCb was sufficient to prevent the generation of apico-basal polarity in 3D-cyst models of epithelial cells. Together, these results further indicate a functional dominance for CLCa *in vivo* and add support to the concept that CLCa plays a housekeeping role in clathrin function that is regulated by competition from CLCb, and that they operate in tandem to modulate clathrin function in epithelial lumen formation.

## Results

### Loss of CLCa reduces post-natal survival and body weight

We have previously shown that CLCa KO and CLCb KO mice have phenotypes in immune cells and neurons (9,22). To further characterise physiological functions of CLCs, heterozygous mice were crossed (*Clta*^+/−^ x *Clta*^+/−^ and *Cltb*^+/−^ x *Cltb*^+/−^) and their offspring compared. When weaned (3 weeks of age), only 9.7% of the *Clta*^+/−^ cross offspring were homozygous for CLCa KO, significantly lower than the expected 25% (p=0.0185), indicating that loss of CLCa reduces survival at the pre- or neo-natal stage (Fig. 1A). In contrast, 23.7% *Cltb*^+/−^ cross offspring were homozygous for CLCb KO, close to the expected 25% (Fig. 1A). Thus, loss of CLCa but not loss of CLCb reduces survival. To identify the developmental stage at which survival is affected, CLCa KO genotypes were analysed pre- and post-natally. At the late embryonic (E18.5) and early post-natal stage (PN1) we observed 30.1% and 27.2% homozygous CLCa KO (*Clta*^−/−^) respectively (Fig. 1B), indicating that loss of CLCa does not affect pre-natal survival. At post-natal day 3 (PN3), significantly fewer than expected mice were homozygous CLCa KO (<10%), which remained the case at PN7 and at 4-weeks old (Fig. 1B) and beyond (data not shown), indicating survival rates are unaffected by the loss of CLCa after PN3. Thus, CLCa has a critical role in post-natal mouse development and/or survival behaviour, such as response to critical feeding cues, within the first 3 days of birth.

**Figure 1.**
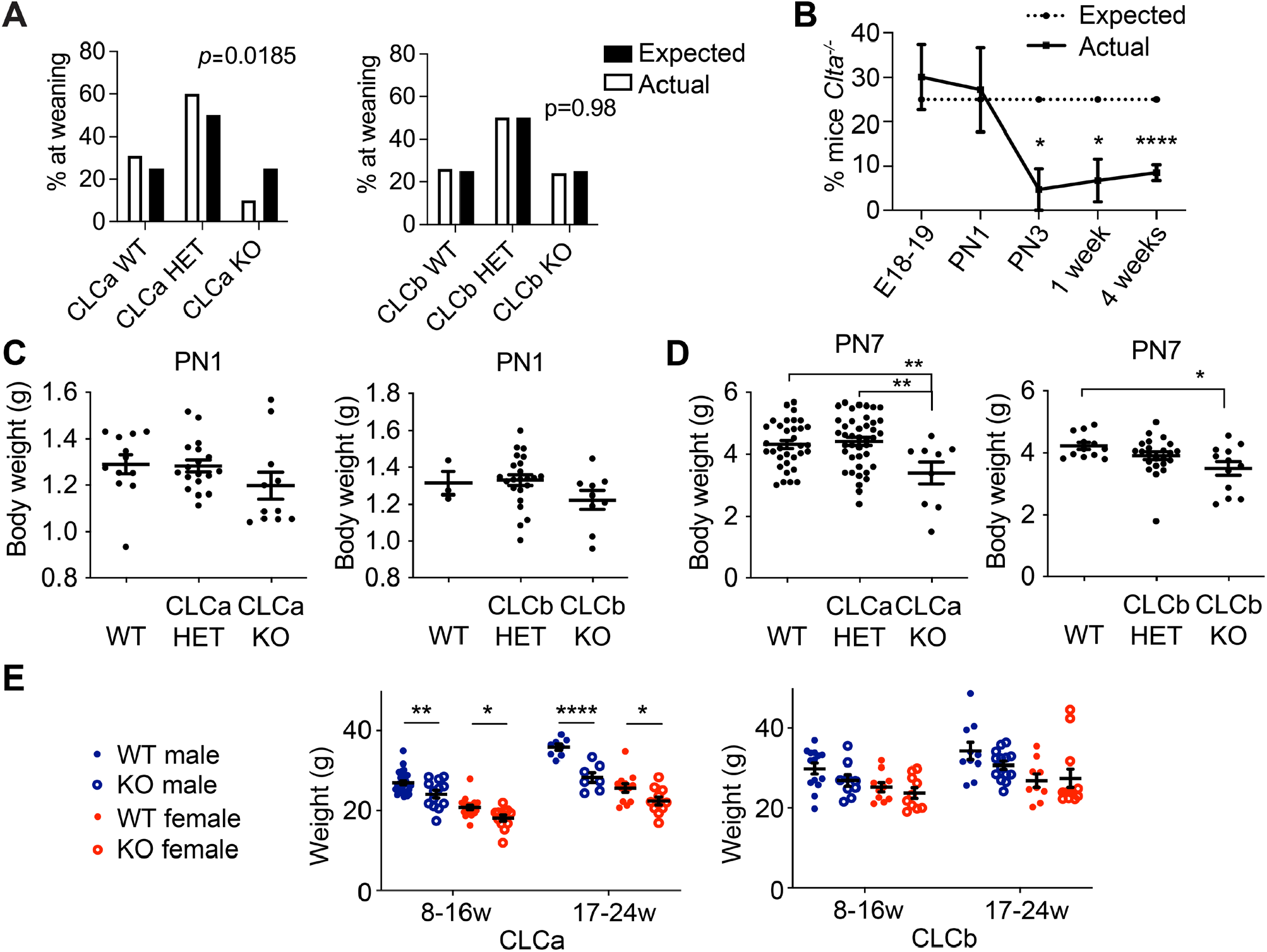
Survival and body weight of mice lacking genes encoding CLCa and CLCb. (A) Breeding cages with *Clta*^+/−^ x *Clta*^+/−^ or *Cltb*^+/−^ x *Cltb*^+/−^ parental genotypes were established and the genotype of offspring analysed at weaning (3 weeks old). Expected and observed percentages for the genotypes *Clta*^+/+^ (CLCa WT), *Clta*^+/−^ (CLCa HET), *Clta*^−/−^ (CLCa KO), *Cltb* ^+/+^ (CLCb WT), *Cltb* ^+/−^ (CLCb HET), *Cltb* ^−/−^ (CLCb KO) are shown. Number of mice analysed: CLCa WT = 303, CLCa HET = 589, CLCa KO = 96, total = 988; CLCb WT = 181, CLCb HET = 344, CLCb KO = 163, total = 688. p values generated by Chi Square analysis comparing the observed genotype percentage of the mice with the expected Mendelian ratios. (B) Genotype analysis of CLCa KO mice at developmental stages following *Clta*^+/−^ x *Clta*^+/−^ breeding crosses. The percentage of *Clta*^−/−^ mice per litter at E18.5 (n=8 litters), post-natal day 1 (PN1, n= 5 litters), PN3 (n=7 litters), 1 week-PN7 (n=5 litters) and 4 weeks old (n=52 litters) is shown. ^*^p < 0.05, ^****^p < 0.0001, Fisher’s exact test. (C&D) Body weight of mice from *Clta*^+/−^ x *Clta*^+/−^ or *Cltb*^+/−^ x *Cltb*^+/−^ breeding crosses at PN1 (C) and PN7 (D) is shown in grams (g) for the indicated genotypes. Number of mice measured: *Clta*^+/−^ x *Clta*^+/−^ breeding crosses (PN1 =41; PN7 =83) and *Cltb*^+/−^ x *Cltb*^+/−^ breeding crosses (PN1 =35; PN7 =47). *p < 0.05, **p < 0.01, one-way ANOVA test, with Holm-Sidak’s multiple comparison. (E) Body weight of adult male and female mice (aged 8-16 or 17-24 weeks) for wild-type (WT) or homozygous knock-out (KO) genotypes from *Clta*^+/−^ x *Clta*^+/−^ (CLCa) or *Cltb*^+/−^ x *Cltb*^+/−^ (CLCb) breeding crosses. Number of mice measured from *Clta*^+/−^ x *Clta*^+/−^ crosses: Aged 8-16 weeks (CLCa WT male = 30, CLCa KO male = 13, CLCa WT female = 17, CLCa KO female = 12); Aged 17-24 weeks (CLCa WT male = 8, CLCa KO male = 7, CLCa WT female = 12, CLCa KO female = 11). Number of mice from *Cltb*^+/−^ x *Cltb*^+/−^ crosses: Aged 8-16 weeks (CLCb WT male = 14, CLCb KO male = 9, CLCb WT female = 10, CLCb KO female = 10); Aged 17-24 weeks (CLCb WT male = 10, CLCb KO male = 14, CLCb WT female = 9, CLCb KO female =12). *p < 0.05, **p < 0.01, **** p < 0.0001, two-way ANOVA test, with Holm-Sidak’s multiple comparison.

The body weight of surviving CLCa KO and CLCb KO pups at PN1 was not significantly different from that of their respective wild-type (WT) and heterozygous littermates (Fig. 1C). However, at PN7, both CLCa KO and CLCb KO pups had a significantly lower body weight than WT pups (Fig. 1D). Loss of CLCa had a greater effect on pup body weight than loss of CLCb, with an average reduction of 21.52% for CLCa KO animals relative to WT controls at PN7, compared to an average 19.1% weight reduction for CLCb KO pups relative to WT. CLCa KO adult male and female mice (over 8 weeks of age) maintained a significantly lower body weight than WT littermates, with male mice more affected (20.8% reduction in male CLCa KO body weight at 17-24 weeks of age compared to a 12.8% weight reduction for female CLCa KO mice, Fig. 1E). No significant difference in the body weight of CLCb KO mice compared to WT littermates above 8 weeks of age was observed (Fig. 1E).

To establish whether the observed mortality and body weight phenotypes were associated with gross morphological changes in the organs of the CLCa KO and CLCb KO mice, tissue slices from mice of each genotype were analysed by hematoxylin and eosin staining. No abnormalities in the structure or morphology of small intestine, skeletal muscle, kidney and heart tissues from surviving CLCa KO adult mice or CLCb KO adult mice compared to WT adult mice (aged between 5 and 9 months) were observed (Fig. S1).

### Loss of CLCa, but not CLCb, reduces fertility in mice

During routine maintenance of CLCa KO and CLCb KO mouse lines, we observed fewer than expected litters when CLCa KO mice were used for breeding. To analyse this further, breeding cages were set up with combinations of WT, heterozygous and KO animals derived from heterozygous crosses (*Clta*^+/-^ x *Clta*^+/-^ or *Cltb*^+/-^ x *Cltb*^+/-)^, and the number of litters born within 54 days recorded (Fig. 2A). While WT breeding pairs produced 1-2 litters during this period, CLCa KO x CLCa KO breeding crosses produced no offspring at all, suggesting loss of CLCa affects fertility. In contrast, loss of CLCb did not reduce the number of litters born (Fig. 2B). Crosses of heterozygous *Clta*^+/-^ mice with CLCa KO animals also generated a significantly reduced number of litters born within 54 days (Fig. 2B). However, breeding WT with CLCa KO animals in either male/female combination produced litter numbers similar to WT breedings during the 54 day period (Fig. 2A). A separate analysis over a 4 month (120 days) period of breeding showed that when either CLC KO strain was bred with WT, no significant difference in the number of litters or neonates per litter was observed (Fig 2B & 2C). Thus, fertility was reduced between animals with no CLCa or between CLCa KO and animals (male or female) heterozygous for CLCa loss. While this fertility defect was not observed when one mate was WT, these breeding experiments support that loss of CLCa affects fertility.

**Figure 2.**
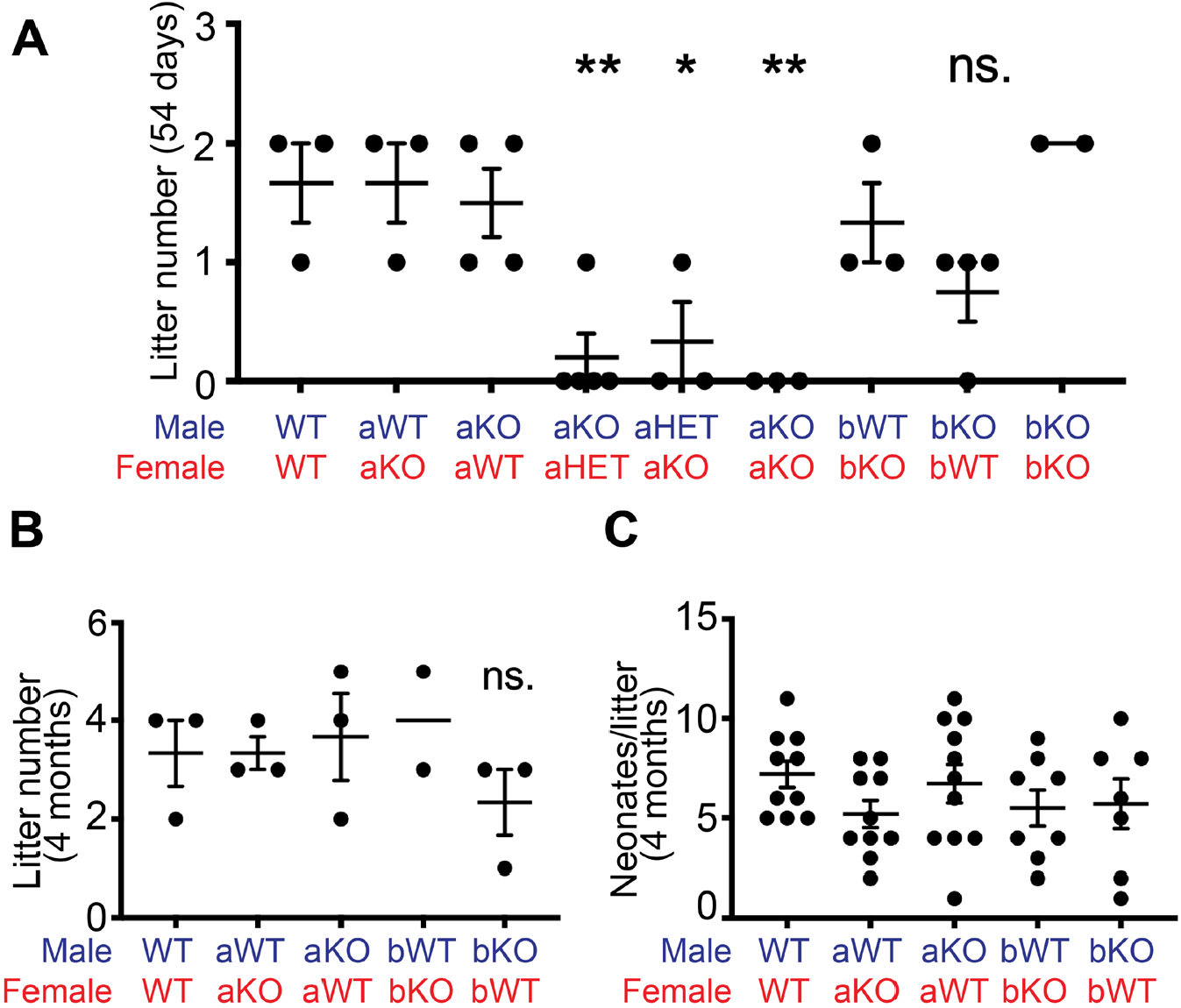
Fertility of mice lacking genes encoding CLCa and CLCb. (A&B) Breeding cages containing male and female mice with the indicated genotypes *Clta*^++^ (aWT), *Clta*^+/-^ (aHET), *Clta*^-/-^ (aKO), *Cltb* ^++^, (bWT), *Cltb* ^+/-^ (bHET), *Cltb* ^-/-^ (bKO) were established and the number of litters born within 54 days (A) or 4 months (B) recorded. Each dot represents the number of litters generated by each breeding pair. Bars represent mean ± SEM, *p < 0.05, **p < 0.01, not significant (ns) compared with WT control, one-way ANOVA test, with Holm-Sidak’s multiple comparison. (C) Breeding cages containing male and female mice with the indicated genotypes were established for 4 months and the number of pups per litter recorded. Graph shows mean ± SEM, one-way ANOVA test, with Holm-Sidak’s multiple comparison.

### Loss of CLCa results in pyometra and reduction in uterine glands

Consistent with loss of CLCa affecting reproduction, we observed that 42.9% of CLCa KO female mice developed a swollen abdomen after 4 months of age due to an enlarged uterus containing cloudy fluid (Fig. 3A & 3B). The uterus is composed of the outer myometrium and the inner endometrium compartments. The endometrium comprises the stroma, glands surrounded by glandular epithelium and the lumen surrounded by lumenal epithelium. H&E staining revealed gross structural abnormalities within the endometrium of CLCa KO mice with enlarged uteri. The cross-sectional width of the endometrium of these mice was more narrow than that in WT animals and the endometrial glandular structures were lost (Fig. 3C & 3D). Quantification confirmed that the number of glands in CLCa KO enlarged uteri was significantly lower than the number of glands in unaffected uteri from other genotypes (WT, CLCb KO and CLCa KO) (Fig. 3D).

**Figure 3.**
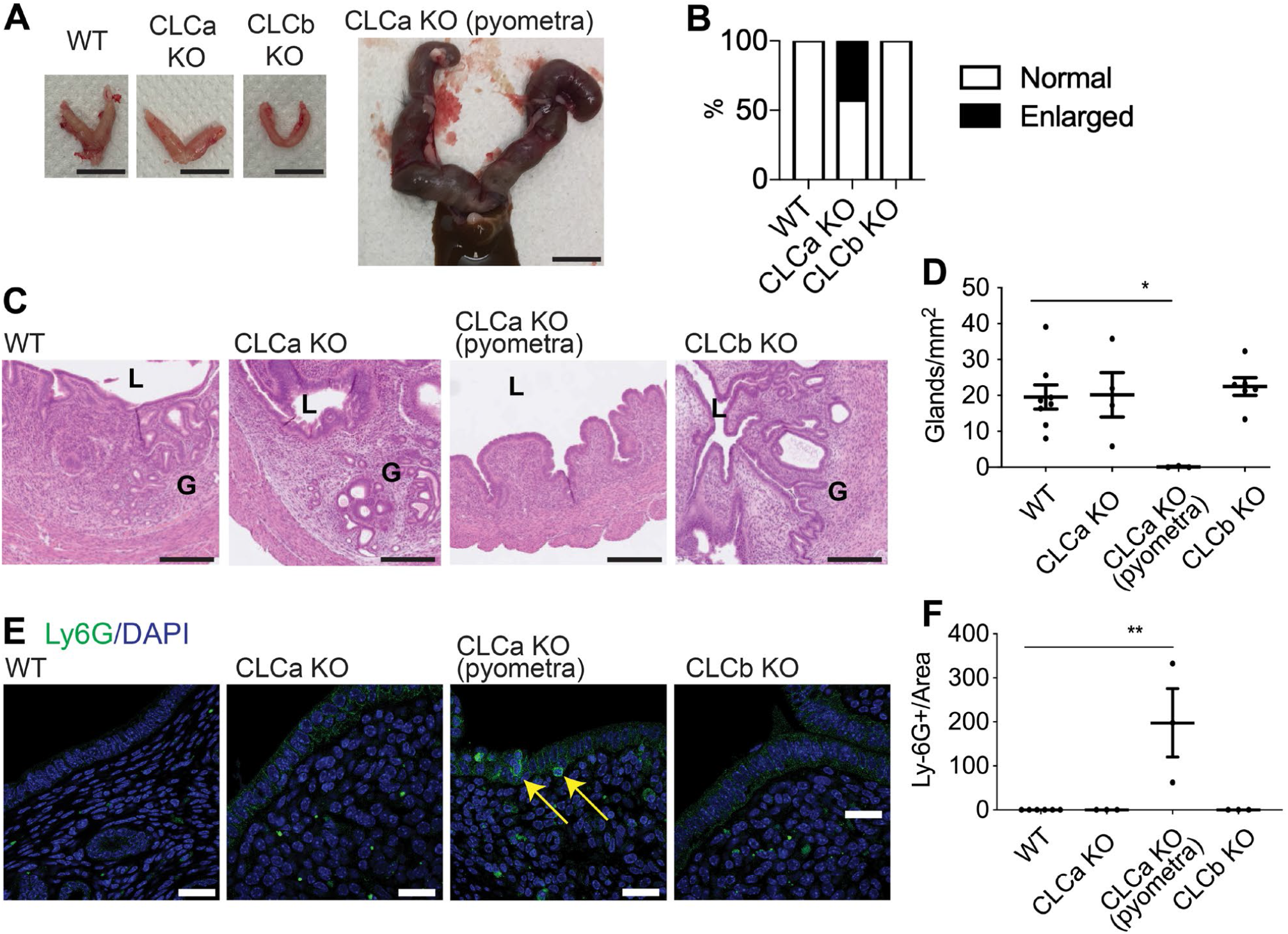
CLCa KO female mice develop pyometra. (A) Images of the uterus from WT, CLCa KO (unaffected) and CLCb KO mice and a CLCa KO mouse with pyometra. Scale bar = 1 cm. (B) The percentage of WT, CLCa KO and CLCb KO female mice that developed an enlarged uterus after 4 months of age. n=31 per genotype. (C) H&E staining of the uterus from WT, CLCa KO and CLCb KO mice and a CLCa KO mouse with pyometra. Images shown are the representative crossed-sectioned images of at least 3 mice in each genotype or condition. L = lumen, G = glands. Scale bar = 250 μm. (D) Number of glands per area (mm2) of uterine tissue cross-section from WT, CLCa KO and CLCb KO mice and a CLCa KO mouse with pyometra from (C). Graph shows mean ± SEM. Number of mice analysed (one section per animal): WT= 8 ; CLCa KO (unaffected) = 4 ; CLCa KO (pyometra) = 3; CLCb KO = 6. *p < 0.05, one-way ANOVA with Holm-Sidak’s multiple comparison. (E) Slices of uterine tissue of the indicated genotype were fixed and stained with antibody against the Ly6G neutrophil marker (of inflammation). Arrows indicate Ly6G-positive cells. Nuclei were stained with DAPI (blue). Scale bar = 25 μm. (F) Quantification of the number of Ly6G-expressing cells per mm^2^ tissue from (E). Each dot represents the average number of Ly6G expressing cells in endometrial tissue of 1 mouse. At least 2 confocal images were counted per mouse. Graph displays mean ± SEM; Number of mice analysed: WT=6; CLCa KO (unaffected) = 3; CLCa KO (pyometra) = 3; CLCb KO = 3. **p < 0.01, one-way ANOVA test, with Holm-Sidak’s multiple comparison.

An enlarged uterus filled with cloudy fluid can be indicative of pyometra, a uterine infection characterised by inflammation and a ‘pus-filled’ uterus (24). To determine whether CLCa KO mice with an enlarged uterus had developed pyometra, the inflammation status of the uteri of these mice was examined and compared to unaffected uteri from other genotypes (WT, CLCb KO and CLCa KO). When sections of uterus were stained with antibodies against the neutrophil marker Ly6G, an indicator of inflammation (25). Ly6G-positive cells were detected in the uterine epithelial layer of CLCa KO mice with an enlarged uterus but not in unaffected animals of any genotype (Fig. 3E & 3F). This sign of neutrophil infiltration is consistent with pyometra as the cause of the enlarged uterus in CLCa KO mice.

To further understand the role of the CLCs in the uterus, we investigated the expression levels of CLCa and CLCb in the uterus. Relative levels of CLCa vs CLCb were determined for whole tissue lysates from the uterus, spleen and brain of WT, CLCa KO (without pyometra) and CLCb KO mice by immunoblotting using the antibody CON.1, recognising the 22 amino acid CLC consensus sequence shared between CLCa and CLCb (26). Thus, the protein levels of the different CLCs can be directly compared within a blot. In the uterus of WT mice, CLCa protein levels were marginally higher than CLCb levels, with CLCa comprising 60.5% ± 3.5% of total CLCs, similar to their relative expression in liver (9) and is not as extreme as CLCa dominance in spleen (Fig. 4A). More balanced levels of the two CLC isoforms were observed for nCLCa and nCLCb in brain (Fig. 4A) and CLCa vs CLCb in muscle (9). Note that the apparent dominance of nCLCb in brain tissue is due to the presence of cells with non-neuronal CLCa that co-migrates with nCLCb during SDS-PAGE (as detected by CON.1 in the CLCb KO mice). Only a slight apparent increase in CLCb level was observed in the tissues of the CLCa KO mice, and vice versa, as noted in analysis of other tissues from these mice (9) (Fig. 4A). Thus development of pyometra correlates with CLCa loss and inability of residual or slightly elevated CLCb to support the clathrin function needed for the impaired pathway, revealing differences in the functions supported by the two CLCs.

**Figure 4.**
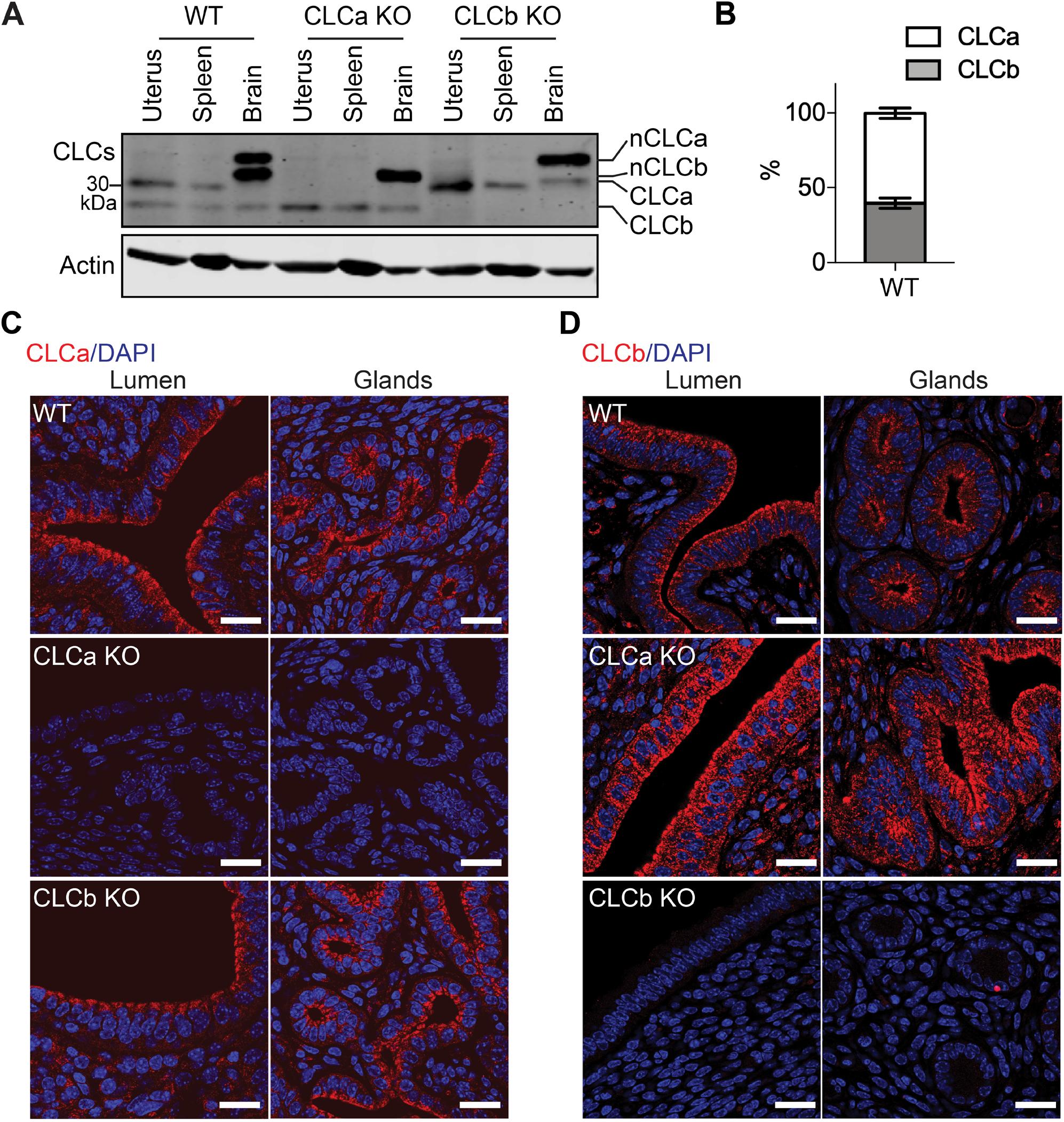
Expression and localisation of CLCa and CLCb in the uterus. (A) Tissue lysates (50 mg of protein) from the uterus, spleen and brain of WT, CLCa KO or CLCb KO mice were analysed by SDS-PAGE and the levels of CLC isoforms compared by immunoblotting using the antibody CON.1 that recognises the consensus sequence shared by CLCa and CLCb and their neuronal splice variants nCLCa and nCLCb. Migration position of molecular mass marker in kilodaltons (kDa) is shown left. (B) Quantification of the amount of CLCa and CLCb found in the in the uterus of WT mice by immunoblotting, shown as a percentage of the total CLC level. n=3. (C&D). Slices of uterine tissue of the indicated genotype were fixed and stained with antibodies against CLCa (C) or CLCb (D), both shown in red. Nuclei were stained with DAPI (blue). Representative images of endometrial lumen and glands are shown. Scale bar = 25 μm.

The relative localisations of CLCa and CLCb in the uterus of WT, CLCa KO and CLCb KO mice was then determined by immunofluorescence. Both CLCa and CLCb were highly enriched in the lumenal and glandular epithelia of the uterus, with both concentrated at the apical pole of the epithelial cells (Fig. 4B & 4C). This is consistent with a role for CLCa at apical domain of epithelia, previously suggested by the specific binding of the epithelial splice variant of myosin VI to CLCa (21) and suggests an additional role for CLCb.

### CLCa loss alters endometrial epithelial cell identity

The advent of pyometra suggests a defect in epithelial integrity, leading to inflammation. To further assess the consequences of CLCa loss on endometrial epithelia, we analysed the levels of FOXA2 expression in the uterus of WT, CLCa and CLCb KO mice. FOXA2 is a transcription factor that regulates epithelial differentiation and development, and its loss affects gland development, uterine function and fertility (27,28,29). In the mature endometrium, expression of FOXA2 is confined to glandular epithelium and absent from epithelia bordering the uterine lumen, as seen the uterine tissue from WT mice (Fig. 5A & 5B). Ectopic expression of FOXA2 was observed in epithelial cells at the uterine lumen of CLCa KO animals with pyometra and also, to a lesser extent, in the CLCa KO mice that did not suffer from pyometra (Fig. 5A and 5B). In contrast, loss of CLCb did not change FOXA2 expression in the uterus, with expression seen only in the glandular epithelium (Fig. 5A & 5B). FOXA2 has previously been shown to regulate the proliferation of endometrial epithelia (30,31). To assess whether ectopic expression of FOXA2 in the lumenal epithelia of the uterus of CLCa KO mice altered cell proliferation, we analysed the expression of the postulated endometrial stem cell marker SRY-Box transcription factor 9 (SOX9) (32) and the proliferation marker Ki67. However, the expression of both SOX9 and Ki67 in endometrial epithelial cells was unaffected by the loss of CLCa or CLCb compared to WT animals (Fig. S2A-C), indicating that the ectopic expression of FOXA2 in the endometrial epithelia does not affect proliferation of these cells. The endometrial epithelium normally consists of simple columnar epithelial cells, however overexpression of FOXA2 in the endometrium has previously been shown to induce epithelial stratification (33). We therefore analysed the expression of keratin 14 (K14), a marker of stratified squamous epithelial cells (34). As expected, the endometrial epithelia of WT, CLCb KO and CLCa KO mice without pyometra did not express K14 (Fig. 6A and 6B). In contrast, K14-expressing cells were found in endometrial epithelia in uteri cross-sections from 40% (2 of 5 analysed) CLCa KO mice with pyometra (Fig. 6A and 6B). Consistently, the cells expressing K14 had a more squamous shape (Fig. 6A), indicating a change in epithelial subtype in the uterus of some of the mice with pyometra.

**Figure 5.**
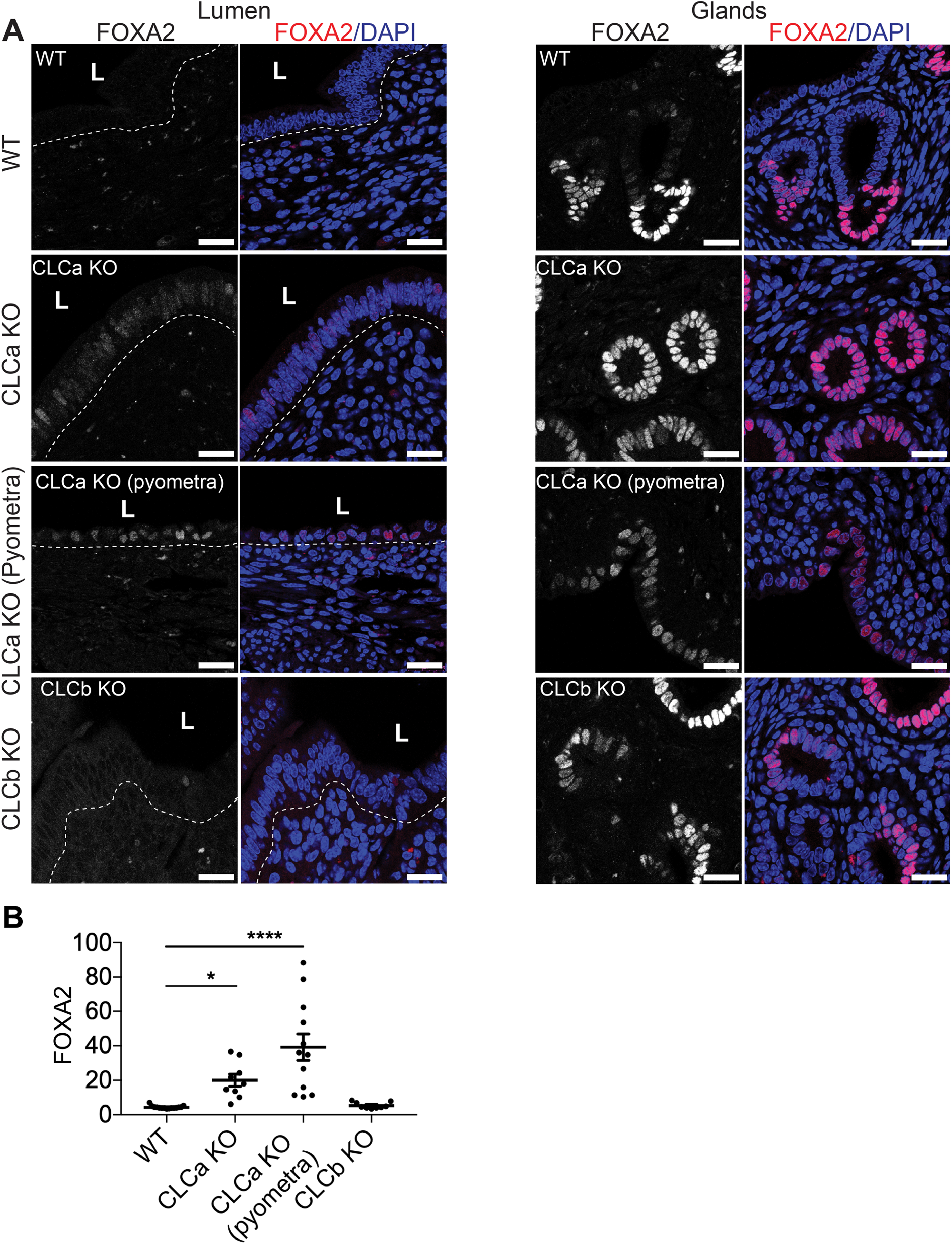
FOXA2 is ectopically expressed in lumenal cells of the endometrium epithelium in CLCa KO mice. (A) Immunostaining for FOXA2 in endometrium from WT, CLCa KO (+ pyometra) or CLCb KO mice. Slices of uterine tissue of the indicated genotype and phenotype were fixed and stained with antibodies against FOXA2 (pink in merge). Nuclei were stained with DAPI (blue). The dotted line shows the boundary between the lumenal epithelium and the rest of the endometrium and the position of the lumen (L) is indicated. Representative images of endometrial lumen and glands are shown. Scale bar = 25 μm. (B) Mean fluorescence intensity for FOXA2 in cells of the lumenal epithelium. Each dot represents the average mean fluorescence intensity of the masked nuclear region of all lumenal cells in one confocal image as assessed by Image J. Three images were analysed per animal. Graph displays mean ± SEM; Number of mice analysed: WT=5; CLCa KO (unaffected) = 3; CLCa KO (pyometra) = 4; CLCb KO = 3. *p<0.05, ****p < 0.0001 on One-way ANOVA test, with Holm-Sidak’s multiple comparison.

**Figure 6.**
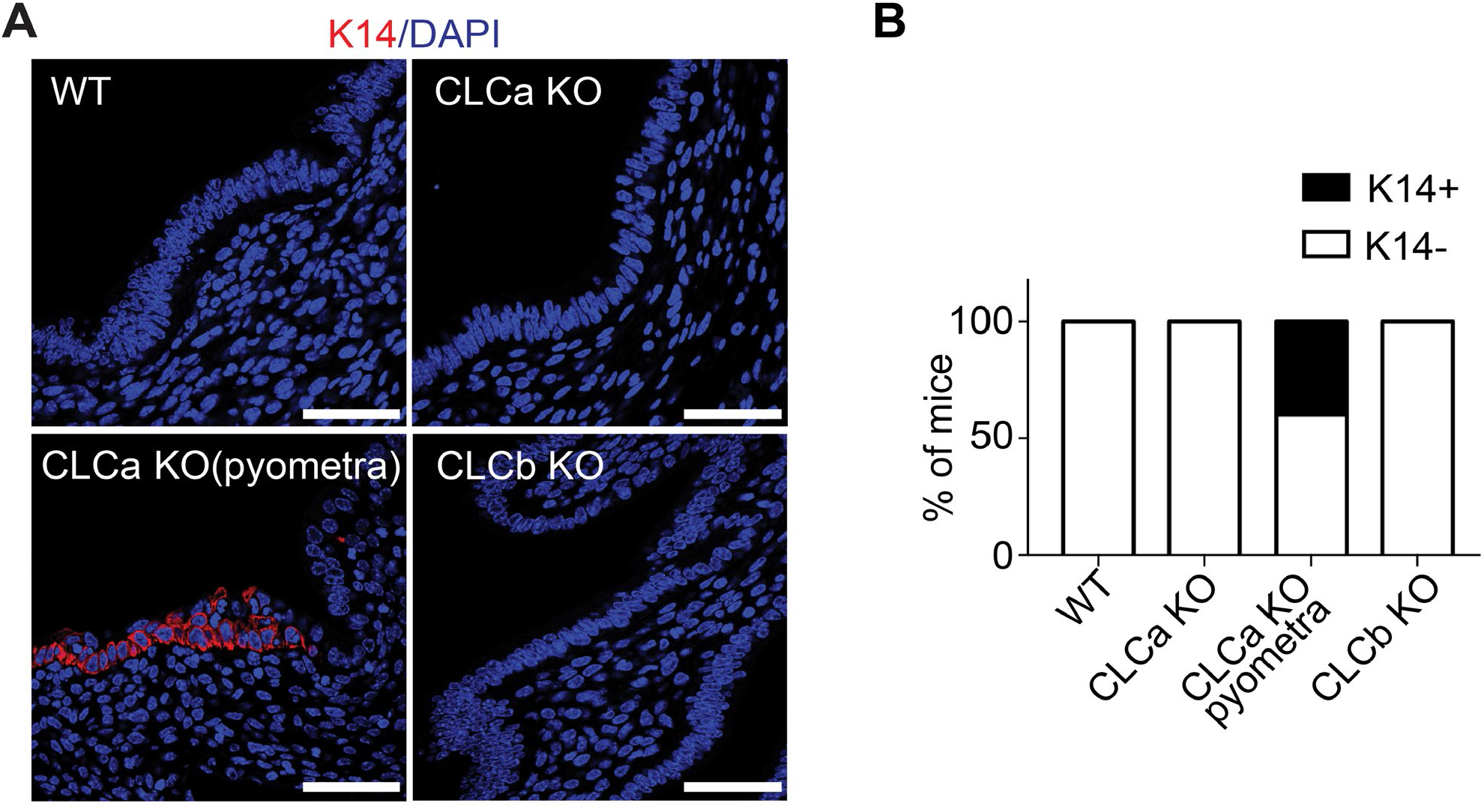
The stratified epithelial marker K14 is ectopically expressed in the endometrium of a subset of CLCa KO mice with pyometra. (A) Immunostaining for K14 in the endometrium of WT, CLCa KO or CLCb KO mice. Slices of uterine tissue of the indicated genotype were fixed and stained with antibody against K14 (red). Nuclei were stained with DAPI (blue). Scale bar = 25 μm. (B) Quantification of the percentage of mice for each genotype that had uterine lumenal epithelial cells expressing K14 (K14+) or did not have any uterine epithelial cells expressing K14 (K14-) as determined by immunostaining. One section per mouse. Number of mice analysed: WT=8; CLCa KO = 4; CLCa KO (pyometra) = 5; CLCb KO = 5.

The structure and integrity of the endometrial epithelia at the lumen in the CLC KO mice was further assessed by immunofluorescent staining of epithelial adherens junction component E-cadherin and the apical tight junction protein ZO-1. WT, CLCa KO with and without pyometra and CLCb KO mice all displayed normal distribution of both E-cadherin and ZO-1, with ZO-1 distributed at the apical surface of the endometrial epithelium and E-cadherin distributed at the basolateral membrane (Fig. S3A & S3B). These findings suggest that epithelial junctions are formed normally in all CLC KO mice. Thus, at the resolution detectable by confocal microscopy, loss of CLCa or CLCb does not alter the organisation or polarity of the endometrial epithelia. The expression and distribution of mucin 1 (MUC1) was also examined, since MUC1 is found at the apical surface of uterine epithelia and acts as a barrier against microbial invasion (35). Loss of this barrier could therefore lead to infection and pyometra. However, a layer of MUC1 was observed at the apical surface of endometrial epithelia of the uterine lumen from WT, CLCa KO and CLCb KO animals (Fig. S3C), suggesting that a loss of apical MUC1 is not the cause of pyometra in CLCa KO animals. Thus the pyometra phenotype may be linked to regional disruption of the epithelium by aberrant conversion to stratified epithelial cells, this differentiation defect being the only defect detected in our analysis.

### Loss of CLCa or CLCb disrupts epithelia cyst polarity *in vitro*

Analysis of uterine tissue suggested that CLCs play a role in epithelial cell differentiation. However, despite the high level of expression of CLCa and CLCb in these cells, loss of either CLCa or CLCb did not result in observable organisational defects in the epithelial layer of the uterine lumen of CLCa KO or CLCb KO mice. The mice used in this study are a constitutive knock-out model, and it is possible that independent compensatory pathways or some substitute functions shared by CLCa and CLCb may have been activated and are sufficient to obscure individual functions of each CLC isoform in these cells. To address this possibility, we assessed the ability of cultured epithelial cell lines to form 3-dimensional cysts *in vitro* upon acute knockdown of either CLC or clathrin heavy chain (CHC17). Ishikawa cells (human endometrial adenocarcinoma cells) and Caco-2 cells (human colorectal adenocarcinoma cells) were treated with control siRNA or siRNA targeting CLCa, CLCb or CHC17 for 72h. Treated cells were then trypsinised and seeded as single cells embedded within the extracellular matrix substrate Matrigel® and grown for 4-6 days to allow cysts to form prior to analysis by immunofluorescence. Ishikawa and Caco-2 cells treated with control siRNA formed symmetrical, spherical cysts with a single lumen surrounded by an F-actin-rich apical membrane (Fig. 7A and 7B). In both Ishikawa and Caco2 cysts, CLCa and CLCb displayed distinct distribution patterns relative to each other (Fig. 4C and 4D). In the cysts, CLCa had a more ubiquitous distribution while CLCb localisation was enriched towards the apical surface of the cells (Fig. 7A and 7B). The 3D-cysts formed by Ishikawa and Caco-2 cells treated with control siRNA displayed the expected basolateral distribution of E-cadherin and ZO-1 localised tightly around the actin-rich lumen (Fig. 7C). A complete loss of this apico-basal polarity was seen in Ishikawa or Caco-2 cysts depleted of CLCa or CHC17 and for Caco-2 cysts depleted of CLCb. In all of these cases, neither E-cadherin or ZO-1 were polarised (Fig. 7C-E), with no symmetrical distribution of cells around a lumen (Fig 7C-E). Analysis of E-cadherin, ZO-1 and actin distribution in Ishikawa cysts depleted of CLCb showed that some apico-basal polarity is obtained in these cysts, with E-cadherin showing a degree of basolateral distribution, and greater symmetry observed across the cyst than for CLCa-depleted cysts (Fig.7C). When cyst size was quantified, Caco2 cysts were found to be reduced in size when depleted for CLCa, CLCb or CHC17, while this was not observed consistently for Ishikawa cysts treated in the same way (Fig. 7F and 7G). However, the cellular distribution in CLC-depleted Ishikawa cysts was not as organised as in the control-treated cysts, and we frequently observed an abnormal number of lumens within the cysts (Fig.7C). During *in vitro* cyst growth, a single lumen develops next to the forming apical surface at the centre of the cysts (36), and therefore an abnormal lumen number is an indicator of defective apico-basal polarity in cysts. To quantify this, a luminogenesis assay was performed. Cells were treated with siRNAs prior to growing cysts for 6 days. For the final 24h, cysts were treated with cholera toxin to expand the lumen, then fixed, and the number of lumens scored according to the phenotypes shown for Caco2 cysts in Fig. 7H. While approximately 80% of control treated cysts contained the expected single lumen for both Caco-2 cysts and Ishikawa cysts (Fig. 7I-J), this was significantly reduced in all cases by treatment with siRNA targeting CLCa. Treatment with siRNA targeting CLCa also significantly increased the proportion of cysts that contained no lumens, while treatment with siRNA targeting CLCb increased the proportion of cysts with multiple lumens, although this was not statistically significant (Fig. 7I-J), suggesting each CLC contributes to distinct clathrin functions during polarisation. Thus, while constitutive loss of either CLCa or CLCb did not appear to disturb the apico-basal polarity of the endometrial epithelia in the knock-out mice, acute knockdown of either CLCa or CLCb was sufficient to disturb the apico-basal polarity of 3D cysts from endometrial and colonic epithelial cells *in vitro*, suggesting the CLCs play an important role in epithelial cell development. The more severe phenotype seen for Ishikawa cysts upon CLCa siRNA treatment compared to CLCb siRNA treatment suggests that CLCa plays a dominant role in epithelial cell development in this tissue.

**Figure 7.**
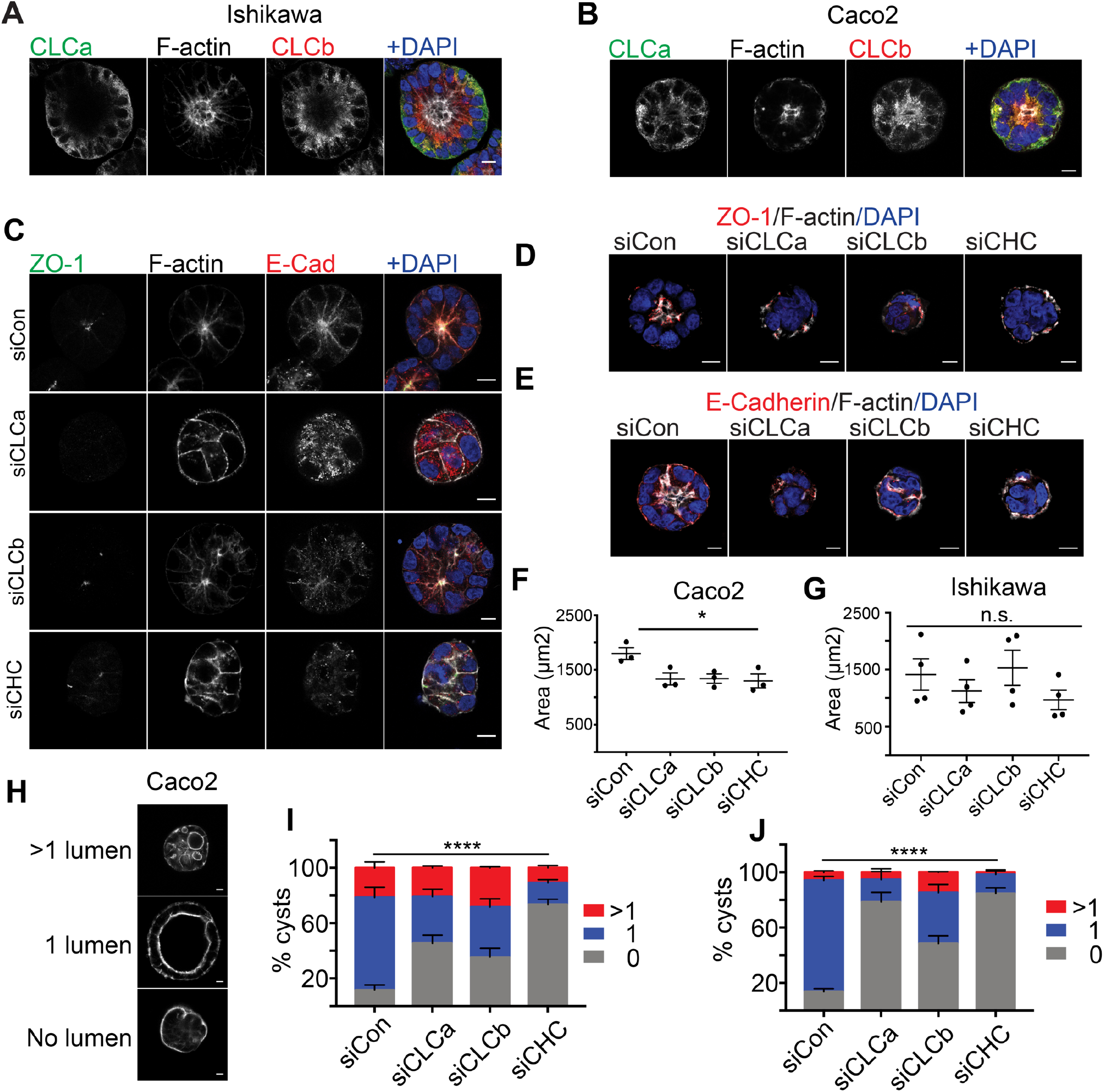
Acute deletion of CLCa or CLCb in Ishikawa and Caco2 cells disrupts epithelial cyst formation and polarisation. Ishikawa (A) or Caco2 (B) cells were seeded as single cells in Matrigel® and grown for 6 days to allow cyst formation. Cysts were treated with cholera toxin for the final 24h to expand the lumens for visualisation. Cysts were fixed and immunostained with antibodies against CLCa (green in merge) and CLCb (red in merge). F-actin and nuclei were stained with phalloidin (grey) and DAPI (blue) respectively. Scale bar 10 μm. Ishikawa (C) or Caco2 (D&E) cells were transfected with siRNA to deplete CLCa (siCLCa), CLCb (siCLCb) or CHC17 (siCHC), or with control siRNA (siCon). 72h after siRNA transfection, cells were trypsinised and seeded in as single cells in Matrigel® and grown for 4 days to allow cyst formation. Cysts were fixed and immunostained with antibodies against ZO-1 (green in (C), red in (D)) and E-Cadherin (red in C&E merge). F-actin and nuclei were stained with phalloidin (grey) and DAPI (blue) respectively. Scale bar = 10 μm. Ishikawa (F) or Caco2 (G) cells were transfected with siRNA to deplete CLCa (siCLCa), CLCb (siCLCb) or CHC17 (siCHC), or with control siRNA (siCon). 72h after siRNA transfection, cells were trypsinised and seeded in as single cells in Matrigel® and grown for 4 days to allow cyst formation prior to fixation. The cross-sectional area of cysts was visualised with antibodies against E-cadherin and the F-actin stain phalloidin and measured using ImageJ software. Each dot represents the average size of over 100 cysts from a single experiment. * p=0.0317, one-way ANOVA. Number of experiments performed: Ishikawa cysts = 4; Caco2 cysts = 3. (H-J) Ishikawa or Caco2 cells were transfected with siRNA to deplete CLCa (siCLCa), CLCb (siCLCb) or CHC17 (siCHC), or with control siRNA (siCon). 72h after siRNA transfection, cells were trypsinised and seeded as single cells within Matrigel® and grown for 6 days to allow cyst formation. Cysts were treated with cholera toxin for the final 24h to expand the lumens for visualisation. Cysts were fixed and lumens visualised with the F-actin stain phalloidin. Representative images of Caco2 cysts with multiple lumens, a single lumen or no lumen, Scale bar 10 μm (H). Percentage of Ishikawa cysts with the indicated number of lumens (0, 1, >1). Error bars show = SEM (n=3) (I). Percentage of Caco2 cysts with the indicated number of lumens. Error bars show = SEM (n = 3). ****p>0.0001, two-way ANOVA (J).

## Discussion

Mice lacking the genes encoding clathrin light chain subunits CLCa or CLCb were produced to address how CLC diversity contributes to clathrin function in tissues and the whole organism. Previous studies of these knockout (KO) animals focussed on the most detectable phenotypes displayed in immune cells and neurons (9,22). Recent findings that CLCa (and not CLCb) binds an epithelial-specific splice variant of myosin VI (21) suggested the possibility of epithelial phenotypes, which had not yet been detected in the KO mice. To search for additional phenotypes resulting from loss of each CLC isoform, general properties of the CLC KO mice including survival, fertility and weight were assessed over time and a new phenotype, relating to a uterine epithelial defect, was discovered and characterised. Previously observed mortality (50%) of the CLCa KO mice was established to occur neonatally, and survivors displayed reduced body weight at this stage. While the CLCb KO mice also displayed reduced body weight from this stage, it was less severe and the expected number of homozygous pups survived. CLCa KO mice displayed reduced fertility and uterine pyometra developed in ∼40% of aged female CLCa KO mice (>4 months), while CLCb KO animals had no detectable uterine phenotype. The lumenal uterine epithelium in affected CLCa KO mice showed aberrant expression of the FOXA2 transcription factor. In a subset of mice developing pyometra, this epithelium displayed concomitant expression of K14 keratin, indicating a switch of epithelial phenotype (34,37). The mice with pyometra also had a reduced number of uterine glands, but the glandular epithelium showed normal marker expression. Either CLCa and CLCb depletion prior to *in vitro* cyst formation resulted in aberrant lumen formation for both Ishikawa (uterine) and Caco2 (gut) epithelial cell lines, supporting necessary roles for both CLC isoforms in lumen formation, with more severe defects observed for CLCa depletion. Interestingly, the differential distribution of CLCa compared to CLCb seen in cysts was not observed in mature uterine epithelium, suggesting either that the CLCs play differential roles depending on the pathway of lumen formation (38) or CLCs play differential roles at different stages of epithelial development. Together these findings highlight a requirement for both CLCs in epithelial lumen development and support the previous suggestion from studies of neuronal phenotypes that CLCa sustains key clathrin functions that are fine-tuned by expression of CLCb, such that their balanced function is required for clathrin pathways in different tissues (22).

In our previous analysis of the CLC KO mice, loss of each CLC isoform was shown to differentially affect synaptic vesicle recycling with CLCa KO mice displaying more severe neuronal defects (22). The perinatal mortality and reduced body weight of the CLCa KO mice reported here were also observed in mice lacking endophilin, another regulator of synaptic transmission with similar neurological abnormality (39). Thus, by analogy, the neurological defects could be the critical factor causing high neonatal mortality in CLCa KO mice through a variety of innervation pathways. While neurological defects might indirectly affect fertility, reduced fertility of the CLCa KO mice could be more directly related to the uterine lumen defects observed in the current study. Breeding the CLC KO mice in various combinations showed that CLCa is required for normal fertility. CLCa KO x CLCa KO breeding did not produce offspring, whereas CLCb KO x CLCb KO breeding was normal. Additionally, CLCa HET x CLCa KO (male x female or female x male) breeding showed significantly reduced litter size, though CLCa WT x CLCa KO (male x female or female x male) breeding was unaffected. While more detailed measurements, such as fertilisation, embryo implantation and embryo growth, would be needed to dissect the specific requirement for CLCa, it is tempting to speculate that the fertility defect results from disrupted function of the uterine epithelial lumen in the CLCa KO mice. Implantation involves embryo interaction with the epithelia lumen that is regulated by uterine gland function (40). Defects in both epithelial cell identity and in gland number were observed in CLCa KO animals developing pyometra, which could be the extreme manifestation of uterine epithelial defects over time, since pyometra appeared in aged mice (>4 months).

Pyometra occurs in middle to older-aged small animals (41). In humans, while the incidence of pyometra is relatively rare (0.038 to 0.5% for gynecologic admissions), it is significantly increased in elderly patients (13.6%) (42,43). The causes of pyometra are not entirely defined. Hypothesized causes of pyometra which could be induced by CLC imbalance include age-related changes in epithelial barrier tissue caused by hormonal variation, leading to altered microbial infiltrate and subsequent inflammation (44). Regarding a role for microbial infiltrate, our investigation of epithelial organisation, proliferation and polarity showed no obvious defect in the barrier properties of the uterine endometrium for the CLCa KO animals which did not have pyometra. Those with pyometra did show neutrophil infiltrate, a sign of infection, though mucin expression was not altered. However, due to limited tissue preparation from the affected animals it was not possible to assess thickness of the mucin layer or the presence of antibacterial peptides, which are also associated with uterine epithelial inflammation (45,46). The ectopic expression of FOXA2 in the endometrium of the mice developing pyometra, and their highly reduced gland number, are indicators of endometrial epithelium defects, particularly since glands develop by invagination of the endometrial lumenal epithelium (40). Overexpression of FOXA2 has been shown to induce stratified epithelium in the mouse uterus (33). That all the CLCa KO mice showed exogenous FOXA2 expression in their endometrial epithelium and only some pyometra-affected mice showed the K14 expression phenotype suggests the CLCa KO animals all manifest an epithelial defect that can then be exacerbated by other factors, such as differential hormonal changes, that could induce more severe effects on endometrial identity and subsequent pyometra.

Hormonal changes are linked to pyometra (24,41). Treatment of dogs with progestogens contributes to pyometra and pyometra generally develops during the secretory (luteal) phase of the oestrous cycle where the progesterone levels are elevated and endometrial secretions change, creating an environment that promotes microbiota growth (41,47). Pyometra can also be induced in C57BL/6 mice by estradiol or Bisphenol A (BPA, considered weak oestrogens) (24). The role of sex hormones in uterine epithelial changes and in the development of pyometra could be affected by CLCa loss through alterations in membrane traffic of the GPCRs that respond to these hormones, such as gonadotropin-releasing hormone receptor (GnRH), follicle-stimulating hormone receptor (FSHR) and luteinizing hormone receptor (LHR) (48,49). The identified role of CLCs in the regulation of GPCR trafficking (16,17), particularly in immune cells (9), could also have immunoregulatory effects, which have been linked to development of pyometra. Macrophage infiltration was seen in BPA-treated mice and associated with susceptibility to pyometra (24). Furthermore, mice lacking the immunoregulatory cytokine Metrnl also developed pyometra (50). Given that CLCa expression is dominant in immune cells and altered IgA antibody production in CLCa KO mice is associated with reduced TGFbR2 receptor uptake (9), immune cell defects in CLCa KO mice could also contribute to development of pyometra.

The ectopic expression of FOXA2 in the endometrium of CLCa KO mice and the further expression of K14 in these mice that develop pyometra indicates abnormality of the epithelial lumen. While the K14 expression could result from FOXA2 expression, it is not clear whether these changes in lumenal cell identity are a response to defects in membrane traffic affecting epithelial development or a direct consequence of CLC depletion. Nonetheless, these findings implicate CLCa function in development of the endometrial lumen. This epithelium, in turn, forms glandular epithelium through budding (40). The glandular epithelium appeared normal in our analysis, but gland number was reduced, presumably due to defects in the originating endometrial epithelium. Lumen formation by epithelial cysts *in vitro* depends on formation of the apical membrane initiation site (AMIS) between dividing cells, representing a different mechanism from formation of the uterine lumens (38). While the CLCa and CLCb isoforms have different distributions relative to each other in tissue and cysts, they are both required for lumen formation, suggesting membrane traffic roles for both in establishing polarity. CLCa is enriched at the apical side of the endometrial and glandular epithelia but is present throughout cyst epithelia. In the former, it is only loss of CLCa that affects function, while in the latter both CLCs are required for proper lumen formation. These findings demonstrate that CLC diversity is required for clathrin to function properly during membrane traffic regulating development of epithelia. The specialised but cooperative functions of the CLC isoforms in membrane traffic during epithelial formation mirrors their balanced role in synaptic vesicle traffic, and further supports the view that CLCa plays the dominant roles in influencing physiological homeostasis with necessary regulation by CLCb. This regulation likely results from the fact that CLCs compete with each other for binding to clathrin heavy chain subunits (23) and contribute differential interaction with accessory proteins (21) and kinases, as well as different biophysical properties (22) to clathrin function.

## Methods

### Animals

CLCa or CLCb KO mouse (both C57BL/6 background) were established as previously described (9,22). All mice used in this study (including WT, heterozygotes and knock-outs used for breeding pairs in fertility studies) were produced by breeding *Clta*^*−/+*^ or *Cltb*^−/+^ heterozygotes. For fertility studies, individual breeding pairs of the indicated genotype were housed continuously in a breeding cage, and the litter size and frequency were monitored for 54-days or 4 months as indicated. All animal procedures and breeding were conducted according to the Animals Scientific Procedures Act UK (1986) and in accordance with the ethical standards at University College London.

### Immunoblotting

Mouse tissue was harvested and snap-frozen in liquid nitrogen and stored at -80 °C until further use. Tissue was homogenized in lysis buffer (50mM Tris-HCl, pH7.4, 150mM NaCl, 1% NP-40, 0.5% Sodium deoxycholate, 0.1% SDS, 1mM EDTA) with cOmplete EDTA-free Proteinase Inhibitor Cocktail (Roche) and then incubated on ice for 1 hr. After centrifugation at 10,000 g, 4°C, 10 min, the supernatant was taken and the protein concentration was determined by BCA assay (Thermo Fisher). Proteins were separated by SDS/PAGE gel and transferred to Protran nitrocellulose membrane (0.2 μm, GE Healthcare). Immunoblotting was performed by incubating with primary antibodies diluted in Tris-buffered saline (20mM Tris pH 7.6, 150mM NaCl) containing 2% BSA overnight at 4°C. Primary antibodies used were anti-CLC (CON.1, made in-house) (26) and anti-tubulin (Abcam, 1:10000). Blots were incubated with IRDye800/700-conjugated secondary antibodies (Licor Biosciences, 1:5000), 1 hour, room temperature. Proteins were detected with LiCor Odyssey Imager (Licor Biosciences) and analysed using Image Studio Lite 5.2 software (Licor Biosciences).

### Histology and immunohistochemistry

Mouse uteri were removed and fixed in 4% paraformaldehyde (pH 7.4, Sigma) (PFA) at 4°C overnight. Specimens were dehydrated through graded ethanol solutions, paraffin-embedded and cut into 5 μm sections. Sections were then deparaffinized and hydrated through graded ethanol solutions. For antigen retrieval, sections were incubated in sodium citrate buffer (10 mM sodium citrate, 0.05% Tween 20 [pH 6.0]), EDTA buffer (1 mM EDTA, 0.05% Tween 20 [pH 8]) or Tris-EDTA (10 mM Tris base, 1 mM EDTA, 0.05% Tween 20, [pH 9.0]) for 20 mins at 95 ^o^C. Slides were incubated with blocking buffer 1 (PBS, 2% BSA) for 15 minutes and blocking buffer 2 (PBS, 1%BSA, 0.3% Triton) for 15 minutes before incubation with primary antibodies diluted in blocking buffer 2 at 4°C overnight. Primary antibodies and antigen retrieval method used were anti-CLCa (Sigma, 1:200, citrate), anti-CLCb (Sigma, 1:200, citrate), anti-FOXA2 (Cell Signaling, 1:100, EDTA), anti-ZO-1 (ThermoFisher, 1:100, citrate), E-Cadherin (BD, 1:100, citrate), Ki-67 (Cell Signaling, 1:200, EDTA), anti-SOX9 (Cell Signaling, 1:200, citrate), anti-K14 (Abcam, 1:100, citrate), anti-Mucin 1 (Abcam, 1:200, Tris-EDTA) anti-Ly-6G (1A8)-Alexa 647 (Biolegend, 1:50, citrate), β-catenin-Alexa 488 (BD, 1:50, EDTA). Slides were washed with PBST (PBS, 0.1% Tween-20) and incubated with Alexa Fluor-conjugated secondary antibody (1:500, Thermo Fisher Scientific) and DAPI (1 μg/ml, Thermo Fisher Scientific) for 1 hour. After PBST washes, slides were mounted with Prolong Diamond antifade media (Thermo Fisher Scientific). Images were obtained using a Leica TCS SP8 confocal microscope with a PLAN APO 40X 1.3NA oil immersion objective. Images were captured using Leica LAS X software and subsequently analysed using Image J.

### Cell lines

Ishikawa cells (ECACC) were grown in Minimum Essential Medium (Sigma) supplemented with 5% fetal bovine serum (FBS; Thermo Fisher Scientific), 1% non-essential amino acids (Sigma), 2 mM L-glutamine (Thermo Fisher Scientific) and 100 U/mL penicillin/100 μg/mL streptomycin (P/S) (Thermo Fisher Scientific). Caco-2 cells were grown in Dulbecco’s Modified Eagle Medium (Thermo Fisher Scientific) supplemented with 10% FBS, 2 mM L-glutamine and P/S.

### Cyst formation and siRNA treatment

For cyst formation, Ishikawa and Caco-2 cell lines were transfected with 20 nM siRNA (Qiagen) using JetPRIME (Polyplus Transfection) transfection regent following manufacturers instructions. Target sequences for siRNA used were: CLTA: 5’-AA AGA CAG TTA TGC AGC TAT T-3’, CLTB: 5’-AAG GAA CCA GCG CCA GAG TGA-3’, CLTC: 5’-AAG CAA TGA GCT GTT TGA AGA. Seventy-two hours after transfection, cells were trypsinised into single cells. 6000 cells were mixed with 30% Matrigel (Corning) in PBS and plated in a well of an 8-well EZ Slide (Millipore) which had been pre-coated with 5 ul Matrigel. Cells were incubated in a 37°C, 5% CO2 incubator for 30 minutes to allow Matrigel to set before then being supplemented with 400 μl culture medium. Culture medium was changed every 2 days. Cholera toxin (0.1ug/ml, Sigma) was added 24hr prior fixation where indicated.

### Immunofluorescence staining

Cysts were washed twice with PBS and fixed in 4% PFA at room temperature for 1 hour before unreacted PFA was quenched with glycine. After PBS washes, cysts were permeabilized with 0.1% Triton X-100 in PBS for 30 mins, followed by blocking with blocking buffer (PBS/2% BSA) for 1 hour. Cysts were then incubated with primary antibodies diluted in blocking buffer overnight at 4°C. Primary antibodies used were anti-CLCa (1:250, Sigma), anti-CLCb (1:250, LCB.1, made-in-house (23)), ZO-1 (1:100, ThermoFisher), E-Cadherin (1:100, BD). After PBS washes, cysts were incubated with Alexa Fluor 488 or 647 conjugated-secondary antibody (1:500, Thermo Fisher Scientific) and Alexa-546-conjugated Phalloidin (1:50, Thermo Fisher Scientific) diluted in blocking buffer for 1.5h at room temperature. Cells were washed with PBS and stained using 1 μg/mL DAPI in PBS (Thermo Fisher Scientific) before slides were mounted with Prolong Diamond antifade media (Thermo Fisher Scientific). Images were obtained using a Leica TCS SP8 confocal microscope with a HC PLAN-APO 63X 1.40 NA CS2 oil-immersion objective with LAS X software and subsequently analysed using ImageJ.

### Statistical analysis

Statistical analysis was performed using Prism 7 (GraphPad). Methods of comparison are stated in figure legends. Mean ± standard error is shown.

## Acknowledgements

We acknowledge University College London IQPath, University College London Institute of Neurology, for processing tissue slices and for H&E staining and University College London KLB animal facility for mouse maintenance. This work was supported by Wellcome Trust Grant 107858/Z/15/Z and MRC grant MR/S008144/1 (both to F.M.B).

## Author contributions

YC: Conceptualisation, formal analysis, investigation, methodology, visualisation, writing – original draft, writing – review and editing. KB: Conceptualisation, formal analysis, investigation, methodology, project administration, visualisation, writing – original draft, writing – review and editing. MDC: Formal analysis, investigation, project administration, FMB: Conceptualisation, funding acquisition, project administration, supervision, writing – original draft, writing – review and editing.

## References

1. Briant K, Redlingshöfer L, Brodsky FM (2020) Clathrin’s life beyond 40: Connecting biochemistry with physiology and disease. Curr Opin Cell Biol 65: 141–149. doi:10.1016/j.ceb.2020.06.004

2. Gould GW, Brodsky FM, Bryant NJ (2020) Building GLUT4 vesicles: CHC22 clathrin’s human touch. Trends Cell Biol 30: 705–719. doi:10.1016/j.tcb.2020.05.007

3. Soykan T, Maritzen T, Haucke V (2016) Modes and mechanisms of synaptic vesicle recycling. Curr Opin Neurobiol 39: 17–23. doi:10.1016/j.conb.2016.03.005

4. Brodsky FM (2012) Diversity of clathrin function: New tricks for an old protein. Annu Rev Cell Dev Biol 28: 309–336. doi:10.1146/annurev-cellbio-101011-155716

5. Kaksonen M, Roux A (2018) Mechanisms of clathrin-mediated endocytosis. Nat Rev Mol Cell Biol 19: 313–326. doi:10.1038/nrm.2017.132

6. Wakeham DE, Abi-Rached L, Towler MC, Wilbur JD, Parham P, Brodsky FM (2005) Clathrin heavy and light chain isoforms originated by independent mechanisms of gene duplication during chordate evolution. Proc Natl Acad Sci U S A 102: 7209–7214. doi:10.1073/pnas.0502058102

7. Jackson AP, Seow HF, Holmes N, Drickamer K, Parham P (1987) Clathrin light chains contain brain-specific insertion sequences and a region of homology with intermediate filaments. Nature 326: 154–159. doi:10.1038/326154a0

8. Kirchhausen T, Scarmato P, Harrison SC, Monroe JJ, Chow EP, Chen, Briant, Mattaliano RJ, Ramachandran KL, Smart JE, Ahn AH, Brosius J (1987) Clathrin light chains LCA and LCB are similar, polymorphic, and share repeated heptad motifs. Science 236: 320–324. doi:10.1126/science.3563513

9. Wu S, Majeed SR, Evans TM, Camus MD, Wong NM, Schollmeier Y, Park M, Muppidi JR, Reboldi A, Parham P, et al (2016) Clathrin light chains’ role in selective endocytosis influences antibody isotype switching. Proc Natl Acad Sci U S A 113: 9816–9821. doi:10.1073/pnas.1611189113

10. Ungewickell E, Branton D (1981) Assembly units of clathrin coats. Nature 289: 420–422. doi:10.1038/289420a0

11. Kirchhausen T, Harrison SC (1981) Protein organization in clathrin trimers. Cell 23: 755–761. doi:10.1016/0092-8674(81)90439-6

12. Chen CY, Brodsky FM (2005) Huntingtin-interacting protein 1 (Hip1) and Hip1-related protein (Hip1R) bind the conserved sequence of clathrin light chains and thereby influence clathrin assembly in vitro and actin distribution in vivo. J Biol Chem 280: 6109–6117. doi:10.1074/jbc.M408454200

13. Legendre-Guillemin V, Metzler M, Lemaire JF, Philie J, Gan L, Hayden MR, McPherson PS (2005) Huntingtin interacting protein 1 (HIP1) regulates clathrin assembly through direct binding to the regulatory region of the clathrin light chain. J Biol Chem 280: 6101–6108. doi:10.1074/jbc.M408430200

14. Newpher TM, Idrissi FZ, Geli MI, Lemmon SK (2006) Novel function of clathrin light chain in promoting endocytic vesicle formation. Mol Biol Cell 17: 4343–4352. doi:10.1091/mbc.e06-07-0606

15. Ybe JA, Greene B, Liu SH, Pley U, Parham P, Brodsky FM (1998) Clathrin self-assembly is regulated by three light-chain residues controlling the formation of critical salt bridges. EMBO J 17: 1297–1303. doi:10.1093/emboj/17.5.1297

16. Maib H, Ferreira F, Vassilopoulos S, Smythe E (2018) Cargo regulates clathrin-coated pit invagination via clathrin light chain phosphorylation. Journal of Cell Biology 217: 4253. doi:10.1083/jcb.201805005

17. Ferreira F, Foley M, Cooke A, Cunningham M, Smith G, Woolley R, Henderson G, Kelly E, Mundell S, Smythe E (2012) Endocytosis of g protein-coupled receptors is regulated by clathrin light chain phosphorylation. Curr Biol 22: 1361–1370. doi:10.1016/j.cub.2012.05.034

18. Mukenhirn M, Muraca F, Bucher D, Asberger E, Cappio Barazzone E, Cavalcanti-Adam EA, Boulant S (2021) Role of clathrin light chains in regulating invadopodia formation. Cells 10: doi:10.3390/cells10020451

19. Tsygankova OM, Keen JH (2019) A unique role for clathrin light chain a in cell spreading and migration. J Cell Sci 132: doi:10.1242/jcs.224030

20. Majeed SR, Vasudevan L, Chen CY, Luo Y, Torres JA, Evans TM, Sharkey A, Foraker AB, Wong NM, Esk C, et al (2014) Clathrin light chains are required for the gyrating-clathrin recycling pathway and thereby promote cell migration. Nat Commun 5: 3891. doi:10.1038/ncomms4891

21. Biancospino M, Buel GR, Niño CA, Maspero E, di Perrotolo RS, Raimondi A, Redlingshöfer L, Weber J, Brodsky FM, Walters KJ, et al (2019) Clathrin light chain a drives selective myosin VI recruitment to clathrin-coated pits under membrane tension. Nature Communications 10: 4974. doi:10.1038/s41467-019-12855-6

22. Redlingshöfer L, McLeod F, Chen Y, Camus MD, Burden JJ, Palomer E, Briant K, Dannhauser PN, Salinas PC, Brodsky FM (2020) Clathrin light chain diversity regulates membrane deformation in vitro and synaptic vesicle formation in vivo. Proc Natl Acad Sci U S A 117: 23527–23538. doi:10.1073/pnas.2003662117

23. Brodsky FM, Galloway CJ, Blank GS, Jackson AP, Seow HF, Drickamer K, Parham P (1987) Localization of clathrin light-chain sequences mediating heavy-chain binding and coated vesicle diversity. Nature 326: 203–205. doi:10.1038/326203a0

24. Kendziorski JA, Kendig EL, Gear RB, Belcher SM (2012) Strain specific induction of pyometra and differences in immune responsiveness in mice exposed to 17α-ethinyl estradiol or the endocrine disrupting chemical bisphenol a. Reprod Toxicol 34: 22–30. doi:10.1016/j.reprotox.2012.03.001

25. Lee PY, Wang JX, Parisini E, Dascher CC, Nigrovic PA (2013) Ly6 family proteins in neutrophil biology. J Leukoc Biol 94: 585–594. doi:10.1189/jlb.0113014

26. Nathke IS, Heuser J, Lupas A, Stock J, Turck CW, Brodsky FM (1992) Folding and trimerization of clathrin subunits at the triskelion hub. Cell 68: 899–910. doi:10.1016/0092-8674(92)90033-9

27. Jeong JW, Kwak I, Lee KY, Kim TH, Large MJ, Stewart CL, Kaestner KH, Lydon JP, DeMayo FJ (2010) Foxa2 is essential for mouse endometrial gland development and fertility. Biol Reprod 83: 396–403. doi:10.1095/biolreprod.109.083154

28. Kelleher AM, Peng W, Pru JK, Pru CA, DeMayo FJ, Spencer TE (2017) Forkhead box a2 (FOXA2) is essential for uterine function and fertility. Proc Natl Acad Sci U S A 114: E1018–e1026. doi:10.1073/pnas.1618433114

29. Besnard V, Wert SE, Hull WM, Whitsett JA (2004) Immunohistochemical localization of foxa1 and FOXA2 in mouse embryos and adult tissues. Gene Expr Patterns 5: 193–208. :10.1016/j.modgep.2004.08.006

30. Villacorte M, Suzuki K, Hirasawa A, Ohkawa Y, Suyama M, Maruyama T, Aoki D, Ogino Y, Miyagawa S, Terabayashi T, et al (2013) B-catenin signaling regulates foxa2 expression during endometrial hyperplasia formation. Oncogene 32: 3477–3482. doi:10.1038/onc.2012.376

31. Lin A, Yin J, Cheng C, Yang Z, Yang H (2018) Decreased expression of FOXA2 promotes eutopic endometrial cell proliferation and migration in patients with endometriosis. Reprod Biomed Online 36: 181–187. doi:10.1016/j.rbmo.2017.11.001

32. Tempest N, Maclean A, Hapangama DK (2018) Endometrial stem cell markers: Current concepts and unresolved questions. Int J Mol Sci 19: doi:10.3390/ijms19103240

33. Wang P, Wu SP, Brooks KE, Kelleher AM, Milano-Foster JJ, DeMayo FJ, Spencer TE (2018) Generation of mouse for conditional expression of forkhead box a2. Endocrinology 159: 1897–1909. doi:10.1210/en.2018-00158

34. Moll R, Divo M, Langbein L (2008) The human keratins: Biology and pathology. Histochemistry and Cell Biology 129: 705. :10.1007/s00418-008-0435-6

35. Dharmaraj N, Gendler SJ, Carson DD (2009) Expression of human muc1 during early pregnancy in the human muc1 transgenic mouse model. Biol Reprod 81: 1182–1188. doi:10.1095/biolreprod.109.079418

36. Apodaca G, Gallo LI, Bryant DM (2012) Role of membrane traffic in the generation of epithelial cell asymmetry. Nat Cell Biol 14: 1235–1243. doi:10.1038/ncb2635

37. Nelson WG, Sun TT (1983) The 50- and 58-kdalton keratin classes as molecular markers for stratified squamous epithelia: Cell culture studies. J Cell Biol 97: 244–251. doi:10.1083/jcb.97.1.244

38. Jewett CE, Prekeris R (2018) Insane in the apical membrane: Trafficking events mediating apicobasal epithelial polarity during tube morphogenesis. Traffic: doi:10.1111/tra.12579

39. Milosevic I, Giovedi S, Lou X, Raimondi A, Collesi C, Shen H, Paradise S, O’Toole E, Ferguson S, Cremona O, et al (2011) Recruitment of endophilin to clathrin-coated pit necks is required for efficient vesicle uncoating after fission. Neuron 72: 587–601. doi:10.1016/j.neuron.2011.08.029

40. Kelleher AM, DeMayo FJ, Spencer TE (2019) Uterine glands: Developmental biology and functional roles in pregnancy. Endocr Rev 40: 1424–1445. doi:10.1210/er.2018-00281

41. Hagman R (2018) Pyometra in small animals. Vet Clin North Am Small Anim Pract 48: 639–661. doi:10.1016/j.cvsm.2018.03.001

42. Chan LY, Lau TK, Wong SF, Yuen PM (2001) Pyometra. What is its clinical significance? J Reprod Med 46: 952–956.

43. Akazawa K, Takamori H, Yasuda H (1991) [clinico-statistical study on pyometra in high-aged outpatients]. Nihon Sanka Fujinka Gakkai Zasshi 43: 1539–1545.

44. Baker JM, Chase DM, Herbst-Kralovetz MM (2018) Uterine microbiota: Residents, tourists, or invaders? Front Immunol 9: 208. doi:10.3389/fimmu.2018.00208

45. Johansson ME, Phillipson M, Petersson J, Velcich A, Holm L, Hansson GC (2008) The inner of the two Muc2 mucin-dependent mucus layers in colon is devoid of bacteria. Proc Natl Acad Sci U S A 105: 15064–15069. doi:10.1073/pnas.0803124105

46. Vaishnava S, Yamamoto M, Severson KM, Ruhn KA, Yu X, Koren O, Ley R, Wakeland EK, Hooper LV (2011) The antibacterial lectin regiiigamma promotes the spatial segregation of microbiota and host in the intestine. Science 334: 255–258. doi:10.1126/science.1209791

47. Cox JE (1970) Progestagens in bitches: A review. J Small Anim Pract 11: 759–778. doi:10.1111/j.1748-5827.1970.tb05587.x

48. Sayers N, Hanyaloglu AC (2018) Intracellular follicle-stimulating hormone receptor trafficking and signaling. Front Endocrinol (Lausanne) 9: 653. doi:10.3389/fendo.2018.00653

49. Flanagan CA, Manilall A (2017) Gonadotropin-releasing hormone (gnrh) receptor structure and gnrh binding. Front Endocrinol (Lausanne) 8: 274. doi:10.3389/fendo.2017.00274

50. Ushach I, Arrevillaga-Boni G, Heller GN, Pone E, Hernandez-Ruiz M, Catalan-Dibene J, Hevezi P, Zlotnik A (2018) Meteorin-like/meteorin-β is a novel immunoregulatory cytokine associated with inflammation. J Immunol 201: 3669–3676. doi:10.4049/jimmunol.1800435

