## Supplementary Material for "Clathrin light chains CLCa and CLCb have non-redundant roles in epithelial lumen formation"

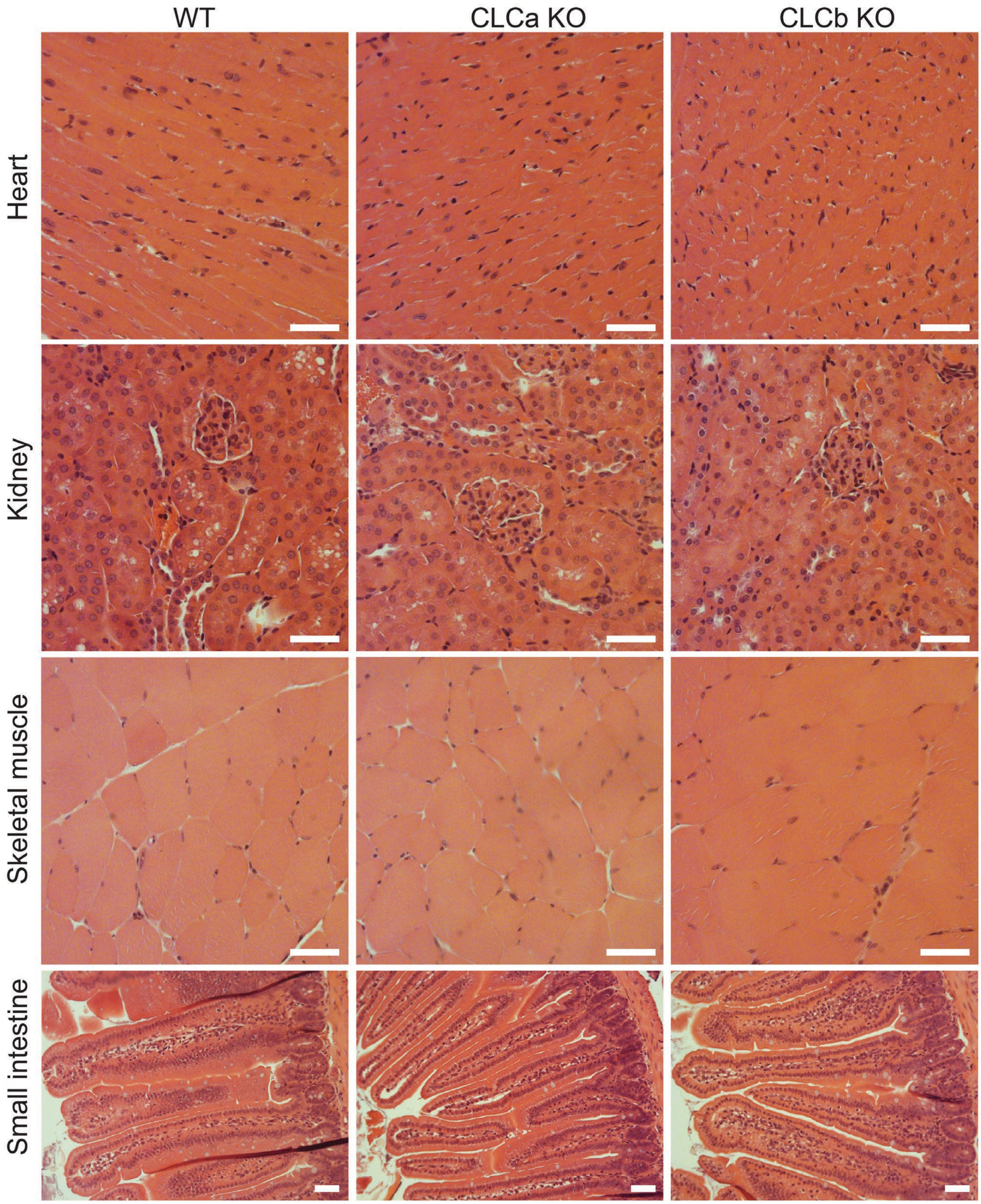


**Figure S1. H&E staining of tissue from CLCa KO, CLCb KO or WT mice**
Tissue from the heart, kidney, skeletal muscle and small intestine of adult WT, CLCa KO or CLCb KO mice was fixed with PFA and stained with H&E. Images shown are the representative bright-field images of at least 3 mice of each genotype. Scale bar = 50 µm.

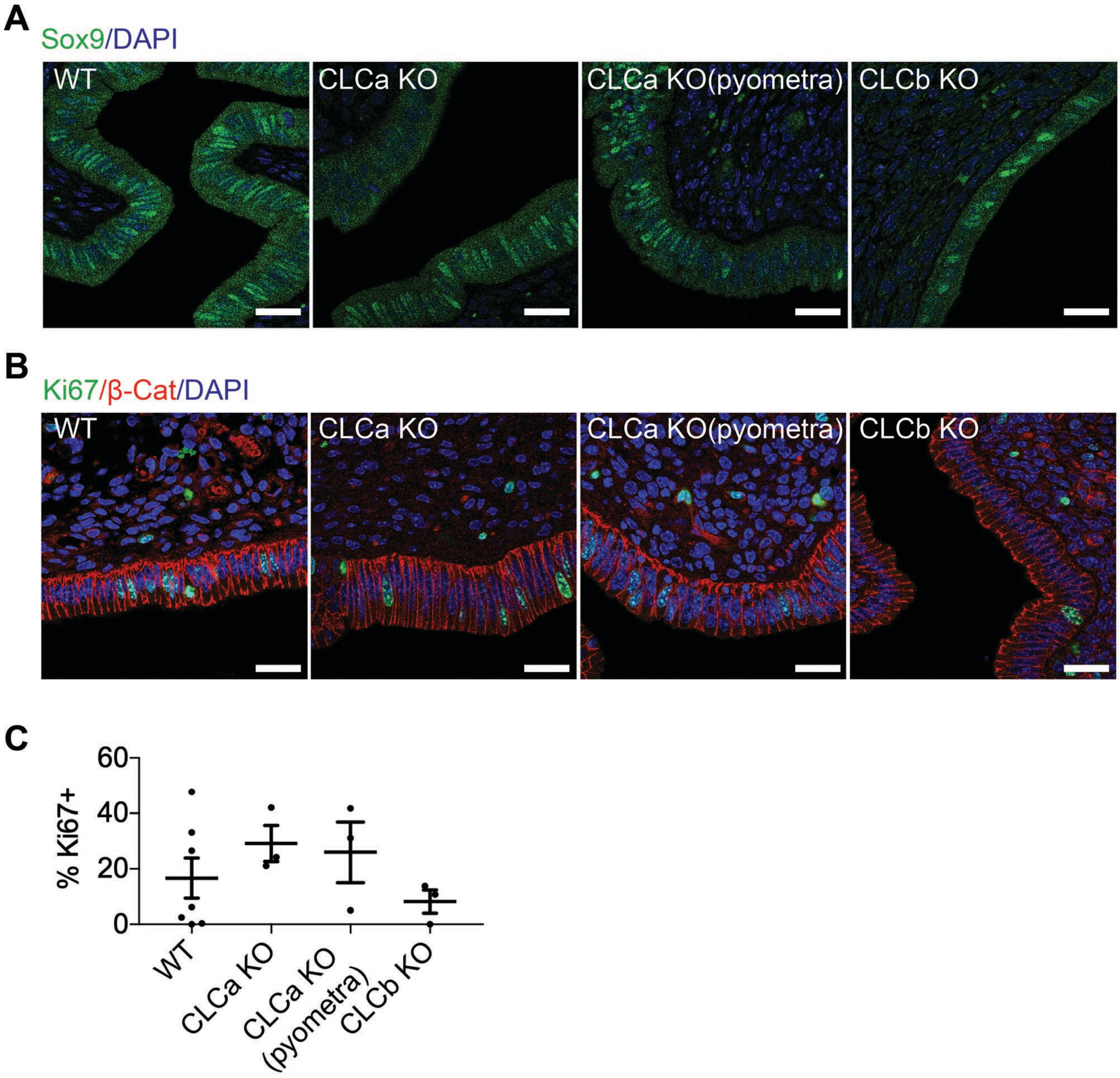


**Figure S2. Expression of SOX9, Ki-67 and b-catenin in CLCa KO or CLCb KO endometrium**
(A) Immunostaining for Sox9 in the endometrium of WT, CLCa KO or CLCb KO mice. Slices of uterine tissue of the indicated genotype were fixed and stained with antibodies against Sox9 (green). Nuclei were stained with DAPI (blue). Scale bar = 25 µm. (B) Immunostaining for Ki67 in the endometrium of WT, CLCa KO or CLCb KO mice. Slices of uterine tissue of the indicated genotype were fixed and stained with antibodies against Ki67 (green) and β-catenin (β-Cat, red). Nuclei were stained with DAPI (blue). Scale bar = 25 µm. (C) Quantification of the percentage of uterine lumenal epithelial cells that express Ki67 as determined by immunostaining. Each dot represents the average percentage of Ki-67 positive endometrial epithelial cells in a single mouse and 2 to 6 confocal images were analysed per mouse. Number of mice analysed: WT=7; CLCa KO = 3; CLCa KO (pyometra) = 3; CLCb KO = 3. Graph shows mean ± SEM. A one-way ANOVA test, with Holm-Sidak's multiple comparison, was performed, with no significant differences between genotypes and phenotypes.

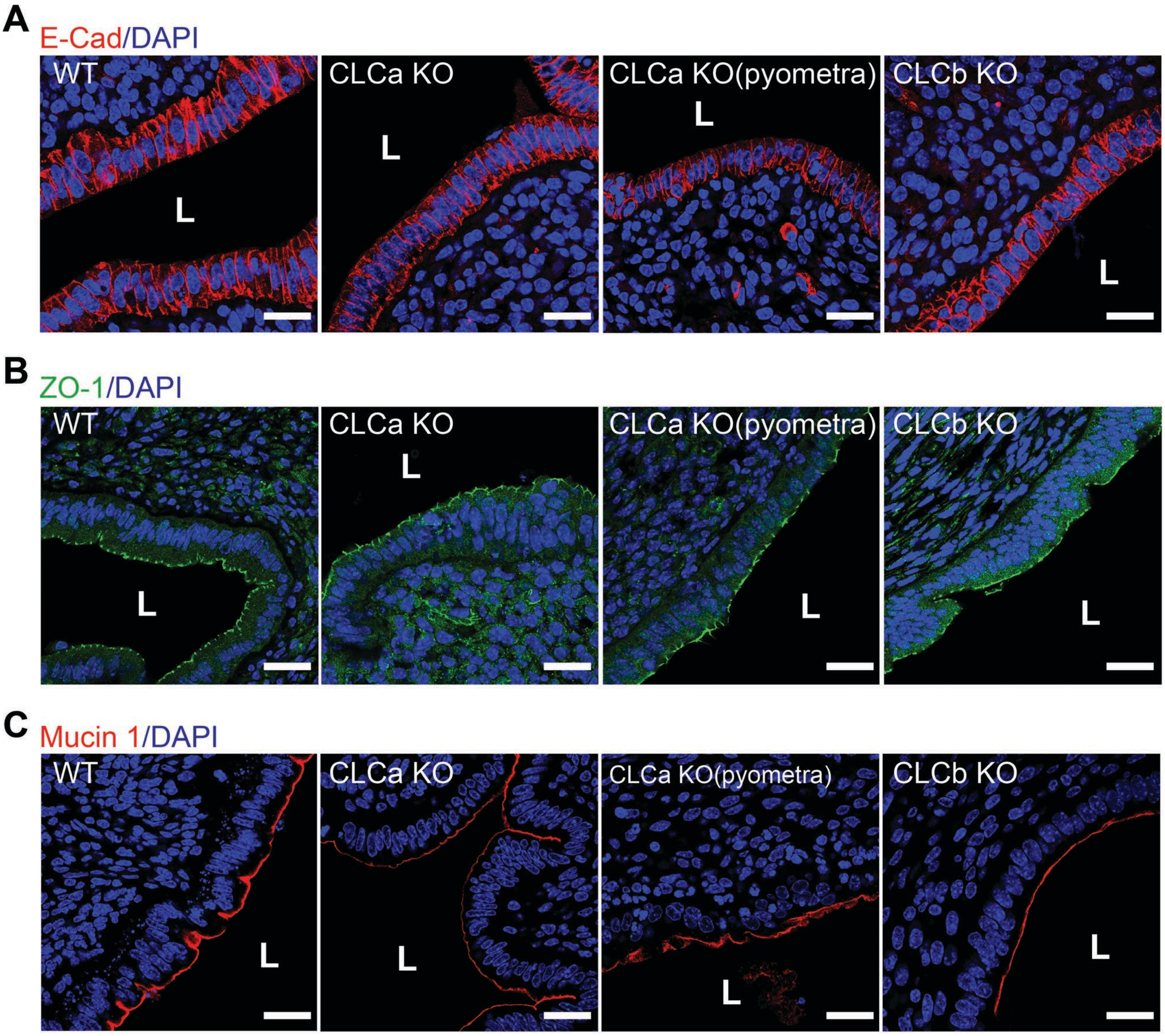


**Figure S3. Expression of E-Cadherin, ZO-1 and Mucin1 in CLCa KO or CLCb KO endometrium**
Immunostaining for E-cadherin, ZO-1, or Mucin 1 in cross-sections of uteri from WT, CLCa KO and CLCb KO female mice. Slices of uterine tissue of the indicated genotype were fixed and stained with antibodies against (A) E-cadherin (E-Cad, red), (B) ZO-1 (green) and (C) Mucin1 (red). Images shown are representative confocal images of at least 2 mice for each genotype. Nuclei were stained with DAPI (blue). Location of the uterine lumen (L) is shown. Scale bar = 25 µm.
